# Multiscale Modeling Identifies Cardiovascular Risk from Common Chemical Exposures

**DOI:** 10.64898/2026.06.09.731239

**Authors:** Shagun Krishna, Xiaoqing Chang, Kristin M. Eccles, Kyle P. Messier, Nicole Kleinstreuer

## Abstract

**Background:** The cardiovascular system is significantly affected by exogenous factors, but understanding the risks posed by pharmaceuticals and environmental chemicals is restricted due to limited data availability. New approach methodologies (NAMs) apply *in vitro, in chemico*, and *in silico* methods to characterize hazard and risk, thus offering rapid, multiscale human biology-based strategies to overcome regulatory challenges and the potential to complement or replace animal testing for understanding chemical cardiovascular effects.

**Methods:** In the present study, we applied a systems-based workflow using physiologically based pharmacokinetic (PBPK) models to convert bioactive concentrations from >300 high-throughput screening (HTS) assays with cardiovascular-relevant molecular and cellular targets to human equivalent administered doses (EADs) for >800 substances with widespread human exposure potential. To derive human-relevant risk predictions, the *in vitro* activity-derived EADs were compared with human exposure estimates and *in vivo* points of departure (PODs) from toxicological animal studies. For a subset of chemicals, we applied a geospatial analysis to assess the combined risks for populations across regions of the US.

**Results:** The combined HTS assay data, human exposure predictions, animal study-based PODs, geospatial exposure data, and PBPK modeling identified compounds with potential cardiovascular toxicity at relevant exposure levels. Personal care product ingredients, flame retardants, herbicides, pesticides, pharmaceuticals, and byproducts of various industrial processes were noted as agents of concern preferentially targeting endothelial cell signaling, nuclear hormone receptors, and other critical cardiovascular targets. Of the 859 chemicals assessed, *in vitro* CV-relevant assays were more risk protective than animal studies for 96.4% of the chemicals. A set of 17 chemicals had a log_10_ bioactivity exposure ratio (BER) below −2, indicating estimated human exposure more than 100-fold above the *in vitro*-derived bioactive dose.

**Conclusions:** This study establishes an integrative, multiscale framework linking molecular perturbations to population-level cardiovascular risk, enabling systematic identification of potentially cardiotoxic chemicals and the communities most vulnerable to their effects. By bridging mechanistic toxicology with pharmacokinetic modeling and epidemiologic context, this approach enhances the biological relevance and translational impact of human health risk assessment. This scalable, adaptable framework supports timely, evidence-based decision-making and aligns with the growing adoption of NAMs to advance cardiovascular research and disease prevention.

**Novelty and Significance:** *What is known?:* High-throughput screening (HTS) assays can identify chemicals with activity at cardiovascular (CV) relevant molecular targets, but translating *in vitro* bioactivity concentrations into biologically meaningful human equivalent doses requires physiologically based pharmacokinetic (PBPK) modelling. The bioactivity exposure ratio (BER) provides a data-driven metric for comparing *in vitro*-derived equivalent administered doses against population exposure estimates, but its application to CV endpoints across a large and chemically diverse environmental chemical landscape has not been demonstrated. Geospatial mapping of CV chemical exposure risk has been demonstrated for a limited set of air pollutants but has not been extended to a broad environmental chemical landscape using human-relevant *in vitro* bioactivity data.

*What new information does this article contribute?:* Integrated *in vitro* to *in vivo* extrapolation (IVIVE) across 859 environmental chemicals demonstrates that cardiovascular-relevant *in vitro* endpoints are sensitive indicators of broader systemic toxicity, 96.4% of chemicals showed positive POD ratios, meaning *in vitro* CV assays flagged hazard at lower doses than non-specific animal toxicity studies despite the absence of endpoint matching. Seventeen chemicals including PFAS, brominated flame retardants, endocrine disruptors, and agricultural herbicides, had a BER below −2, indicating estimated human exposure more than 100-fold above the *in vitro*-derived cardiovascular bioactive dose, with convergent evidence from both *in vitro* and *in vivo* data supporting regulatory priority. County-level geospatial mapping reveals that cardiovascular chemical exposure risk is geographically heterogeneous across the United States, concentrated in industrially active regions already associated with elevated cardiovascular disease mortality, identifying specific populations for targeted environmental monitoring.

*Summary:* This study presents a scalable, systems-based IVIVE framework that integrates cardiovascular-relevant *in vitro* HTS bioactivity data with reverse dosimetry, population exposure predictions, and *in vivo* animal toxicity data to prioritize environmental chemicals for cardiovascular risk assessment. Applied to 859 chemicals spanning personal care products, flame retardants, pesticides, pharmaceuticals, and industrial compounds, the framework demonstrates that CV-relevant *in vitro* endpoints are sensitive indicators of systemic toxicity even in the absence of direct endpoint matching with *in vivo* studies. The BER emerges as a flexible and resource-adaptable prioritization metric, identifying 92 chemicals where estimated human exposure falls within the CV bioactive range, of which 17 represent the highest regulatory priority based on convergent evidence from both data streams. Geospatial mapping further reveals regional heterogeneity in cardiovascular chemical exposure risk concentrated in industrial areas of the central and southeastern United States. This work advances the application of new approach methodologies for cardiovascular chemical risk assessment at a time of accelerating regulatory transition toward human-relevant *in vitro*-based safety evaluation, providing a reproducible computational workflow directly applicable to chemical prioritization under evolving EPA and FDA regulatory frameworks.

## Introduction

Cardiovascular disease (CVD) is a leading cause of mortality and morbidity worldwide, and environmental factors have a substantial impact on CV health outcomes ^1–3^. While genetics and lifestyle factors (e.g., smoking) have well-characterized impacts, up to 23% of CVD can be linked to environmental variables such as air pollution and industrial and agricultural chemicals. However, chemical-specific risks to the cardiovascular system are poorly understood ^4–6^, and cardiotoxic effects are a leading cause of drug attrition, with cardiovascular liabilities often undetected until late-stage clinical trials or post-market surveillance ^7^.

Current cardiotoxicity testing techniques are insufficient to assess the full range of structural, electrophysiological, and contractility related adverse effects, and largely do so independent of the underlying mode of action, accounting for a significant portion of this knowledge gap ^8^. Recent technological and informatics-driven breakthroughs have given rise to “New Approach Methodologies” (NAMs), which have the potential to transform safety evaluations of the continually rising number of environmental and industrial chemicals and significantly enhance drug development pipelines ^9–11^. NAMs are comprised of *in vitro, in chemico*, and *in silico* approaches, or combinations thereof, that are increasingly employed to yield a mechanistic understanding of environmental chemical bioactivity and drug safety, and may be more cost- and time-effective, and human-relevant, substitutes for the use of animals ^12–14^. High throughput screening (HTS) approaches can facilitate preliminary risk evaluation and provide mechanistic insights into toxicity potential ^15–19^. For example, the US EPA has adopted *in vitro* HTS assays and computational models as alternatives for three current Tier 1 assays in the Endocrine Disruptor Screening Program ^12,20–24^ and globally standardized protocols describe NAMs that can substitute an animal test to identify skin sensitizers ^25^. NAMs, which overcome the constraints of conventional animal models are being investigated for the testing and safety evaluation of a range of environmental chemicals and pharmaceuticals for human cardiotoxicity ^11,26–28^. However, the majority of NAMs created for cardiotoxicity have been tailored for use in screening pharmaceutical agents for arrhythmic potential, and a more comprehensive approach is needed to understand the full spectrum of CV effects ^29–31^.

Uncertainty regarding the relationship between *in vitro* dosimetry and *in vivo* exposure levels is a major obstacle to more widespread adoption and predictive utility of such approaches ^32^. Before HTS and other NAM data can be effectively used for risk assessment, drug safety evaluation, novel therapeutic development, and other research and regulatory purposes, it is necessary to analyze the ability of these methodologies to quantitatively inform about biological responses *in vivo*. Various computational/in silico methodologies can be integrated into multiscale systems biology modeling strategies to assist with this contextualization. Computational methods can make predictions about how a chemical will behave in the body based on its structural characteristics, e.g., quantitative structure activity relationship (QSAR) models for predicting absorption and tissue partitioning prediction ^33–36^. Mathematical models include the representation of the pharmacokinetic (PK) behavior of chemicals in the body, covering absorption, distribution, metabolism, and excretion–toxicity (ADMET). These range from simplistic (one-compartment model) to more complex multi-compartment and physiologically based pharmacokinetic (PBPK) models, which are essential in predicting how external exposures relate to internal concentrations ^37^. These models can be paired with other *in silico* and *in vitro* information sources to cover a broad range of chemical and drug exposures and provide both toxicokinetic and toxicodynamic insights ^38–40^.

To understand the risks chemicals may pose to the cardiovascular system, we used CV-relevant in vitro bioactivity assays associated with molecular and cellular events linked to six failure modes of cardiovascular toxicity ^41^ and applied multiscale systems modeling approaches to better contextualize *in vitro* activity values, extrapolate to *in vivo* exposures (IVIVE), and predict geospatial risk across the U.S population. Chemical-specific concentrations representing the most sensitive CV-relevant molecular and cellular perturbations were translated into a range of daily equivalent administered doses (EADs) using multiple IVIVE models under different simulation scenarios and incorporating uncertainty. The derived EADs were further compared with human exposure predictions, geospatial exposure information where available, and *in vivo* point of departure values to develop bioactivity-exposure and POD ratios to provide a risk-based context for identifying compounds of concern. This study presents a novel integrated framework combining NAMs for *in vitro* bioactivity, multiscale systems modeling, and exposure characterization to identify agents that may represent a public health concern via their potential cardiovascular toxicity.

## 2. Methods

### 2.1 Study framework

The overall framework developed and applied in this study is presented in Figure 1. Briefly, *in vitro* HTS data mapped to mechanistic targets linked to CVD were used to identify chemicals with sub-cytotoxic bioactivity concentrations representing critical CV molecular and cellular signaling perturbations, which were converted to EADs using reverse dosimetry and compared with human and animal exposure values. The workflow is briefly described here, and details are provided in Supplementary Methods.

**Figure 1:**
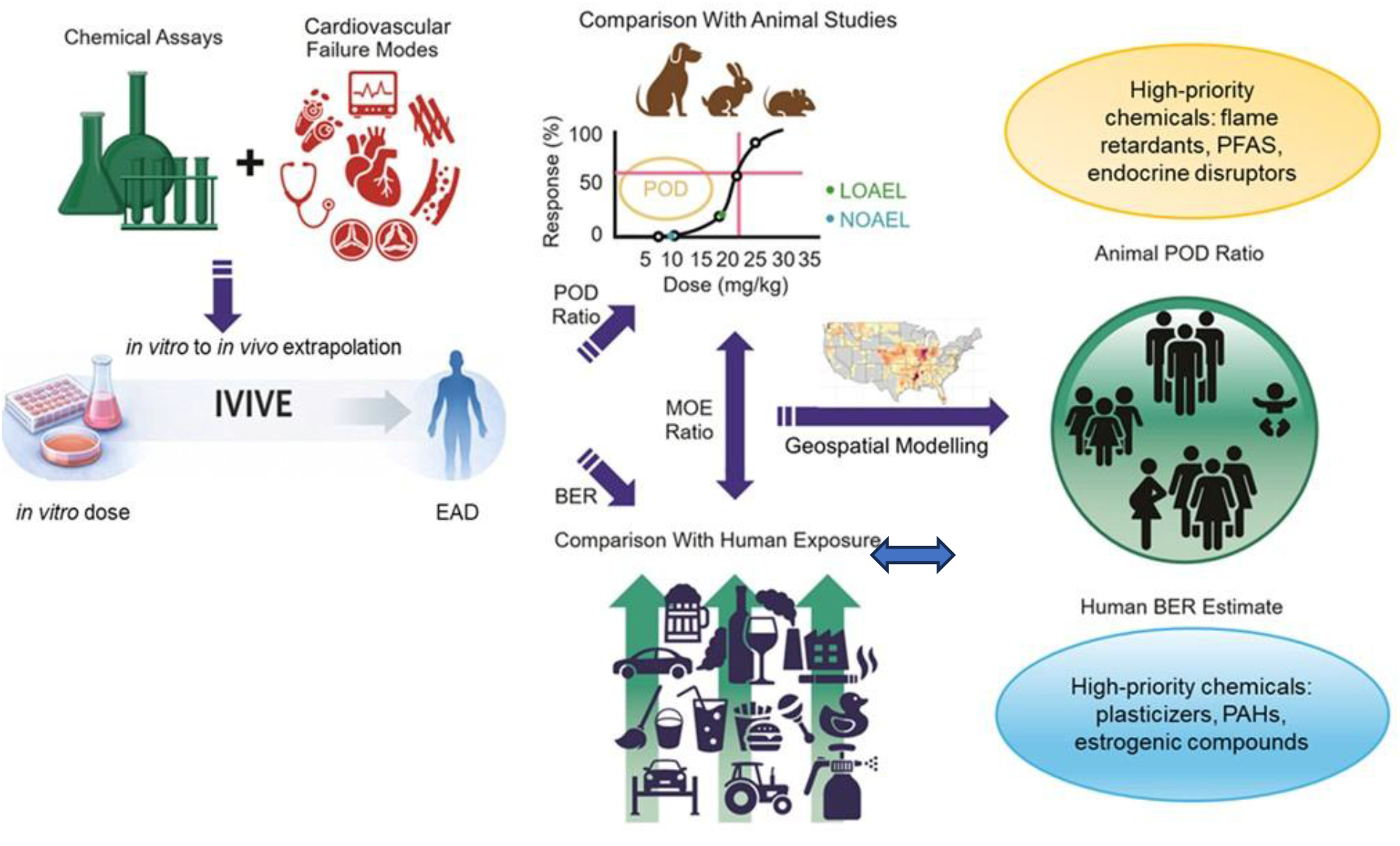
Study workflow. A total of 859 chemicals with *in vitro* HTS bioactivity data mapped to CV targets were processed through multiple IVIVE models to generate a range of equivalent administered doses (EADs). EADs were compared against population exposure estimates (median and 95th percentile) and *in vivo* points of departure (PODs) from toxicological animal studies. Geospatial exposure data were used to create risk maps for the US population.

### 2.2 Curation of *in vitro* bioactivity data

A subset of *in vitro* HTS assays from the US federal Tox21/ToxCast mapped to CV failure modes ^41^ identified chemicals demonstrating specific bioactivity in molecular and cellular targets associated with adverse CV events. Here, we used the most sensitive CV-relevant assay target for each chemical, i.e., the lowest half-maximal activity concentration (MinAC50), following cytotoxicity filtering by applying chemical-specific cytotoxicity thresholds ^42^ as a starting point for the analysis. Uncertainty in bioactivity values was accounted for by deriving upper and lower confidence bounds via statistical bootstrapping using the toxboot R package ^43^.

### 2.3 Calculation of the EAD

The resultant bioactivity data bounds were used to calculate the EADs using the IVIVE tool from the NIH Integrated Chemical Environment (ICE; https://ice.ntp.niehs.nih.gov/) ^44^. IVIVE was applied to calculate EAD values for each chemical that would result in steady-state or maximal plasma concentrations equal to MinAC50s obtained from the CV-relevant *in vitro* assays. Three toxicokinetic models representing the human body were applied (one-compartment (1C), three-compartment (3C), and physiologically-based toxicokinetic (PBTK) model), simulating oral and intravenous (IV) exposure routes under three dosing scenarios with differing duration and dosing interval (3-day 2-hr, 3-day 24-hr, and 30-day 24-hr).

### 2.4 Curation of exposure data

Exposure estimates were derived from the SEEM3 model of the US EPA ExpoCast program ^45,46^. The “US Total Exposure” median and the 95th percentile on the credible interval for the median forecast from the SEEM3 model were available for 815 chemicals in this study ^46^.

### 2.5 Curation of in vivo POD data

The U.S. EPA ToxValDB, accessible via the CompTox Chemicals Dashboard (https://comptox.epa.gov/dashboard/), provided *in vivo* POD data, comprising source-referenced toxicity values gathered from multiple studies. Data were filtered to obtain oral NOAEL, NOEL, LOAEL, and LOEL values in mg/kg-bw/day from mammalian toxicity studies ^47^. For each chemical, we derived the available minimum POD (min-POD_invivo_) value and the p5-POD_invivo_, i.e., the fifth percentile of the POD distributions by applying a discontinuous averaging function in R (quantile).

### 2.6 Calculation of the POD ratio and Bioactivity Exposure Ratio (BER)

To provide exposure- and dose-based context to the *in vitro* bioactivity values, we calculated the POD ratio, the bioactivity exposure ratio (BER), and the margin of exposure (MOE) for each chemical as follows:

log_10_(PODratio) = log_10_(POD_invivo_) - log_10_(EAD_ivive_)

log_10_(BER) = log_10_(EAD_ivive_) - log_10_(EAD_Expocast_)

log_10_(MOE) = log_10_(POD_invivo_) - log_10_(EAD_Expocast_)

### 2.7 Chemical Characterization

Functional use categories were assigned to chemicals with negative BER values using the NIH ICE Chemical Characterization tool ^48^, which draws on EPA’s Chemical and Products Database (CPDat; ^49^.

### 2.8 Geospatial Mapping

Geospatial mapping via the GeoTox framework (Eccles *et al*., 2023, Messier et al. 2025) provides an approach for geospatially mapping *in-vitro* and *in-silico* based dose-response characterization to individual and population exposures. Here, data for potentially cardiotoxic chemicals were obtained from the EPA National Air Toxics Assessment (NATA) Database ^50^, which provides county-level atmospheric concentration estimates for 177 chemicals derived from dispersion and chemical transport models. Only NATA chemicals that were shown to be associated with a CV failure mode, as described above, were retained for this analysis. County-level inhaled doses were converted to internal steady-state plasma concentrations using the PBTK model with county-specific demographics via a Monte Carlo framework (1,000 iterations, Eccles *et al*., 2023). The geographic Exposure Activity Ratio (EAR) was calculated using the median steady-state plasma concentration in a county divided by the most sensitive CV-relevant assay concentration to visualize geospatial CV risk hotspots. Maps were created using the ggplot2 (Wickham, 2009) and sf (Pebesma and others, 2018) R packages.

## 3. Results

### In Vitro Bioactivity Data

This systems-based multiscale framework for predicting chemical impacts on cardiovascular health first leveraged 314 *in vitro* HTS assays mapped to CV failure modes to identify 859 chemicals with potential adverse CV effects ^41^. The chemical-assay bioactivities were refined to focus on sub-cytotoxic concentrations ^42^ and the most sensitive CV-relevant targets extracted for each chemical. The most prevalent failure mode identified was endothelial injury/coagulation (EI/COA), affected by 575 chemicals (Supplementary Figure 3), followed by a change in vasoactivity and cardiomyocyte/myocardial injury. Full details of target frequencies, top assays, failure mode prevalence, and assay platform contributions are provided in Supplementary Figures 1–4.

### Dose Estimation

Three distinct PK models - 1C, Solve 3C, and Solve_PBTK – were applied to conduct IVIVE assessments for both oral and IV exposure routes under three dosing scenarios (3 days 2 Hr, 3 days 24 Hr, and 30 days 24 Hr). EAD_ivive_ results for all model-scenario combinations are provided in Supplementary Table 1. The 1C model consistently predicted higher EAD_ivive_ values than the other models, due to its route-agnostic single-compartment structure. The more complex models, Solve_3C and Solve_PBTK, provided more realistic exposure scenarios and yielded lower EAD_ivive_ values. For oral exposure, chronic dosing (30 days 24 Hr) yielded higher EAD_ivive_ estimates than the multiple-dose scenario (3 days 2 Hr), while IV estimates were similar across scenarios for the 3C and PBTK models. As expected, EAD_ivive_ ranges widened with uncertainty in the *in vitro* AC50 values for all chemicals.

### Dose Comparison

The EAD_ivive_ estimates for different simulation scenarios based on CV-relevant HTS data were compared with the POD_invivo_ (animal toxicity data; 695 chemicals) and EAD_Expocast_ (human exposure predictions; 815 chemicals) and visualized in Figure 2 and are representedon a log10-mg/kg-bw/day basis. Representative results are shown for 3 days 2hr dosing scenario, providing both the most conservative estimates and the most closely aligned with *in vitro* exposure conditions; results for all other scenarios are provided in Supplementary Tables 2 and 3 and Supplementary Figures 5 and 6. Two POD_invivo_ estimates were used the minimum (min-POD_invivo_) and fifth percentile (p5-POD_invivo_) to bound the *in vivo* data (Supplementary Table 2). A positive log_10_PODratio indicates *in vitro* data yielded a lower (more protective) dose than the *in vivo* animal data. A positive log_10_POD ratio indicates that the dose extrapolated from the *in vitro* data was more sensitive than that derived from the *in vivo* animal study. Here, 96.4% of the chemicals had positive log_10_POD ratios, affirming that *in vitro* bioactivities provide a comparable or more protective extrapolated dose than that derived from animal studies. The log_10_POD ratio ranged from −1.63 to 9.7 with similar median values for both measures (3.07 and 3.15 with min-POD_invivo_ and p5-POD_invivo,_ respectively). Five chemicals, Dichlorodiphenyltrichloroethane (DDT), Benzotrichloride, Glutaraldehyde, Urethane, and Fenamiphos, had log_10_POD ratios of less than −1, indicating the dose from the animal study for these compounds was more than an order of magnitude lower than the extrapolated dose from the *in vitro* data. For these chemicals, POD_invivo_ values were derived from chronic or subacute studies reflecting toxicodynamic profiles that unfold over exposure durations exceeding those represented *in vitro*. In 95% of cases (663/695 with both data streams), the human exposure values were below the doses derived from the *in vivo* animal data (five chemicals,perfluorooctanesulfonate (PFOS), perfluorooctanoic acid (PFOA), potassium PFOS, 3-iodo-2-propynyl-N-butylcarbamate, and naphthalene, had negative MOE values, the remaining 27 chemicals could not be evaluated for MOE due to lack of data); however, in a significant fraction of cases (102 chemicals), the human exposure values were above the extrapolated dose from the *in vitro* bioactivity data (log_10_BER < 0; discussed below).

**Figure 2:**
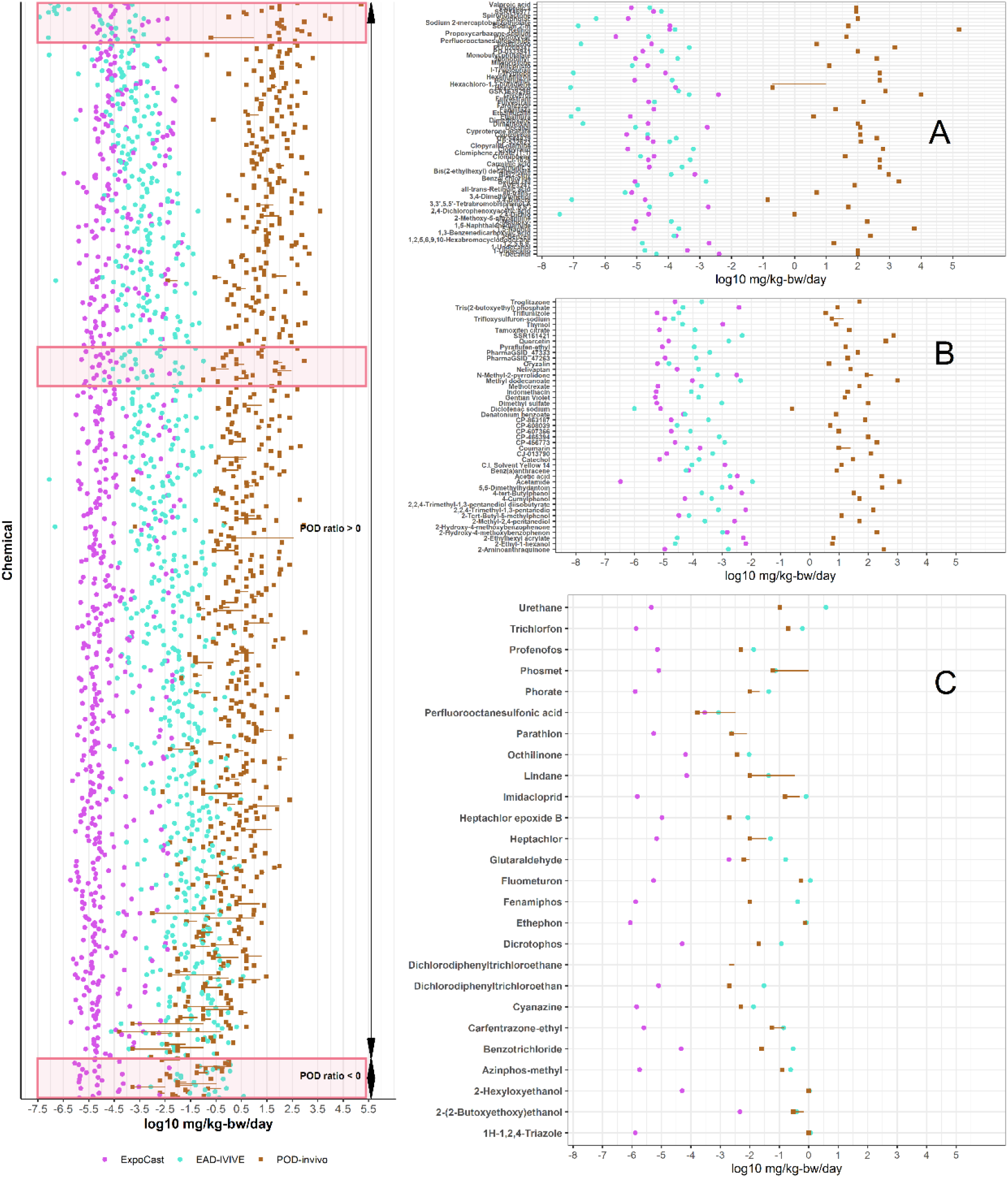
Comparison of Exposure, EAD_ivive_, POD_invivo_, sorted by log10(PODratio) for the 3-day, 2-hr dosing scenario. ExpoCast 95th percentile (pink circles; human exposure estimates), EAD_ivive_ (cyan circles; doses derived from *in vitro* CV-mapped assay data), and POD_invivo_ (brown squares; animal toxicity data spanning log(min-POD_invivo_) to log(p5-POD_invivo_)) are shown. Panels highlight: (A) chemicals with the highest log_10_POD ratios; (B) chemicals near the median log_10_POD ratio; (C) all chemicals with log_10_POD ratio < 0.

The complete results from all simulated dosing scenarios are provided in Supplementary Table 2.

### Exposure Comparison

Human exposure predictions (US total exposure median and 95th percentile, Supplementary Table 3) were compared to EAD_ivive_ estimates under different simulation scenarios using various PBPK models (Figure 3, Supplementary Figures 7 and 8). Figure 3 shows the comparison of EADs and exposure estimates for chemicals with low BER in the multiple dosing scenario (3 days 2 Hr). A total of 102 chemicals had log_10_BER < 0, indicating the potential for exposure to occur within the bioactive dose range extrapolated from *in vitro* data. Among them, 17 chemicals had log_10_BER < −2, meaning the estimated human exposure level was approximately 100-fold higher than the dose associated with in vitro CV perturbations at the molecular/cellular level.

**Figure 3:**
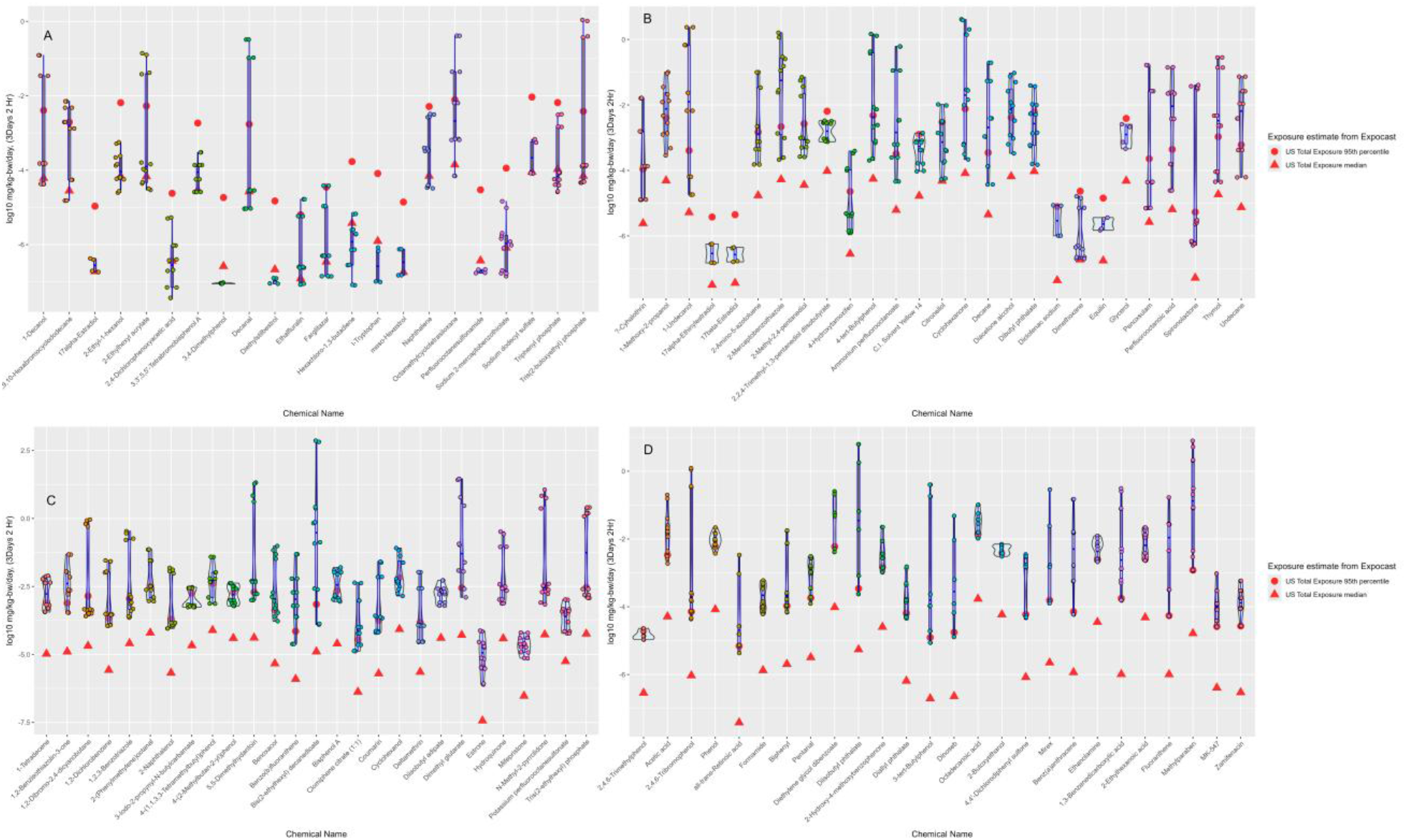
Exposure estimates and EAD_ivive_ values that defined chemicals with log10 BER < 0 for 3 days 2 Hr dosing scenario. (A) represents chemicals with BER < 0 by comparing EAD_ivive_ to both US total exposure median and 95th percentile estimates. Remaining subsections are the chemicals with BER <0 by comparison with 95th percentile exposure estimates.

Numerous chemicals with widespread potential for human exposure, including those used in personal care products, flame retardants, herbicides, pesticides, pharmaceuticals, and various industrial processes, show negative log_10_BER. Functional use categories for all 102 chemicals with negative BER are summarized in Figure 4. Detailed chemical-specific findings and exposure estimates for all three BER tiers are provided in Supplementary Table 3.

**Figure 4:**
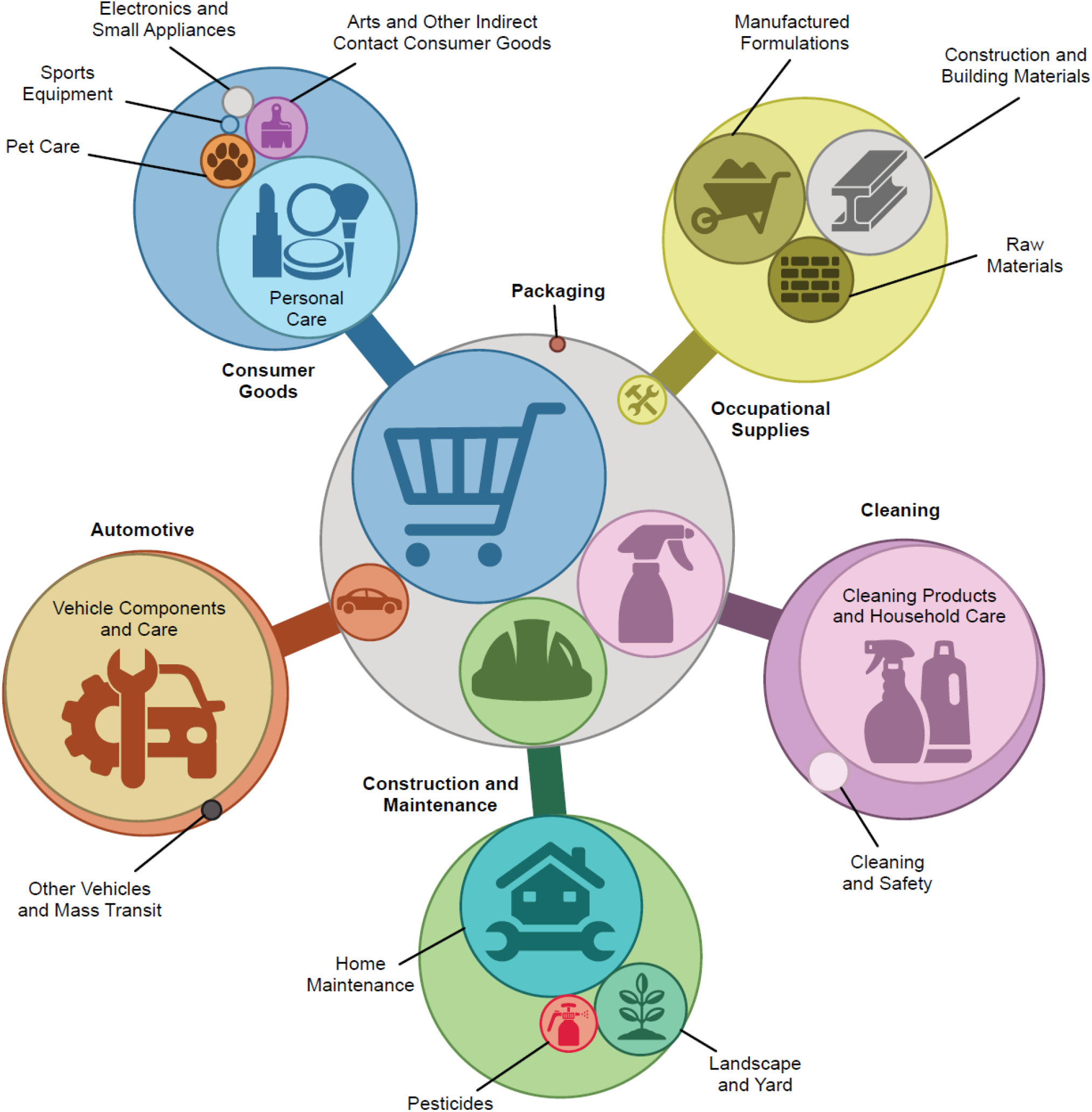
Functional use categories of the 102 chemicals with a negative BER (suggesting that human exposure predictions exceed the extrapolated dose from *in vitro* CV-relevant assays) in the 3-day, 2-hr dosing scenario, as assigned using the ICE Chemical Characterization tool. Categories are organized into six broad sectors: Consumer Goods, Occupational Supplies, Cleaning, Construction and Maintenance, Automotive, and Packaging, with subcategories shown within each sector. A single chemical may be associated with multiple functional use categories.

### Risk characterization across different metrics

To delineate the risk characterization across different metrics, Figure 5 compares sets of chemicals based on Positive and Negative POD ratios, BER, and MOE from the 3 days 2 Hr dosing scenario. Analysis of chemicals with Positive POD ratios (Figure 5 A, where *in vitro* data were more sensitive than *in vivo*) reveals that a majority (n=551) exhibit a positive BER and positive MOE, indicating predicted human exposure is below extrapolated *in vitro* doses and *in vivo* PODs, suggesting an adequate safety margin. Four chemicals (n=4, PFOA, Potassium PFOS, 3-Iodo-2-propynyl-N-butylcarbamate, and Naphthalene) exhibited both a negative BER and a negative MOE, indicating that human exposure exceeds thresholds derived from both *in vitro* and *in vivo* data, warranting further investigation and risk characterization. Interestingly, no chemicals displayed both a negative BER and a positive MOE, underscoring the internal consistency of the two metrics within this group.

**Figure 5:**
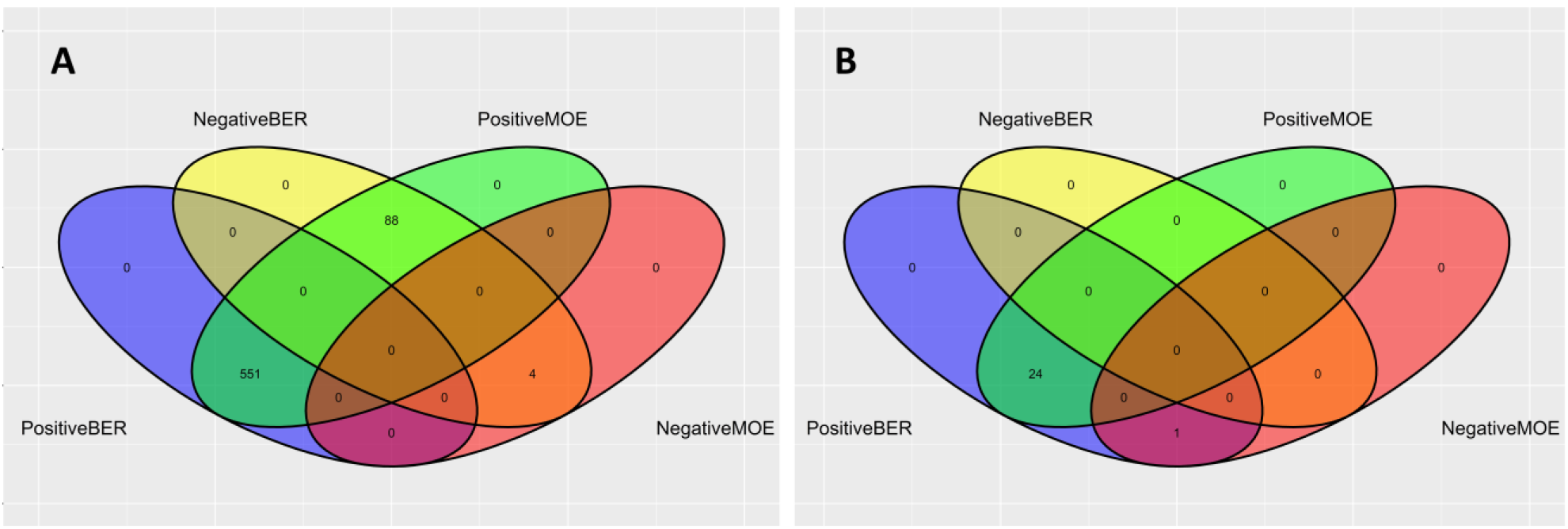
Chemical distributions based on POD ratios, BER, and MOE from a 3-day, 2-hour dosing simulation. Panel A presents chemicals with positive POD ratios (*in vitro* more sensitive than *in vivo*), and Panel B focuses on chemicals with negative POD ratios (*in vivo* more sensitive than *in vitro*) in 3 days 2hr dosing scenario. Positive BER (human exposure lower than *in vitro* extrapolated dose), Negative BER (human exposure higher than *in vitro* extrapolated dose), Positive MOE (human exposure lower than *in vivo* dose), Negative MOE (human exposure higher than *in vivo* dose).

Furthermore, for chemicals with negative POD ratios (Fig 5B), in which *in vivo* studies were more sensitive than the *in vitro* assays, all 25 chemicals exhibited a positive BER, indicating that human exposure remains below the *in vitro*-derived threshold. Of these, 24 chemicals also had a positive MOE, confirming that exposure does not exceed *in vivo*-derived safety levels either. A single chemical, PFOS had a positive BER but negative MOE, indicating that while human exposure was predicted below the *in vitro*-derived bioactive dose, it marginally exceeds the *in vivo*-derived POD, warranting further attention. Notably, no chemical in this group exhibited a negative BER, confirming that even when *in vitro* assays were less sensitive than *in vivo* studies, estimated human exposure did not surpass the *in vitro*-derived bioactive dose in any case. These comparisons can help inform the identification of chemicals for priority risk assessment and potential regulatory action.

### Geospatial Mapping

Two NATA chemicals, hydroquinone and parathion, had activity in the vasoactivity assays, the EAR for all counties of the USA did not exceed 0.001 and thus are not presented in Figure 6. Ten chemicals showed activity in the endothelial injury and coagulation assays, and of those, three chemicals had an EAR>0.001 and are presented in Figure 6. Two out of three chemicals that had activity in the cardiomyocyte/myocardial injury assays had EAR>0.001. Hexachloro-1,3, butadiene is both widespread and had high EAR values for the endothelial injury and coagulation failure mode, with a few counties showing EAR>0.001 for the cardiomyocyte/myocardial injury failure mode. Naphthalene exposure was also widespread, affecting endothelial injury and coagulation and cardiomyocyte/myocardial injury failure modes, however, it does not exceed EAR values of 0.002. Also informing on the endothelial injury and coagulation failure mode, 2,4-Dichlorophenoxyacetic acid only had a few counties that were above the EAR threshold of 0.001.

**Figure 6:**
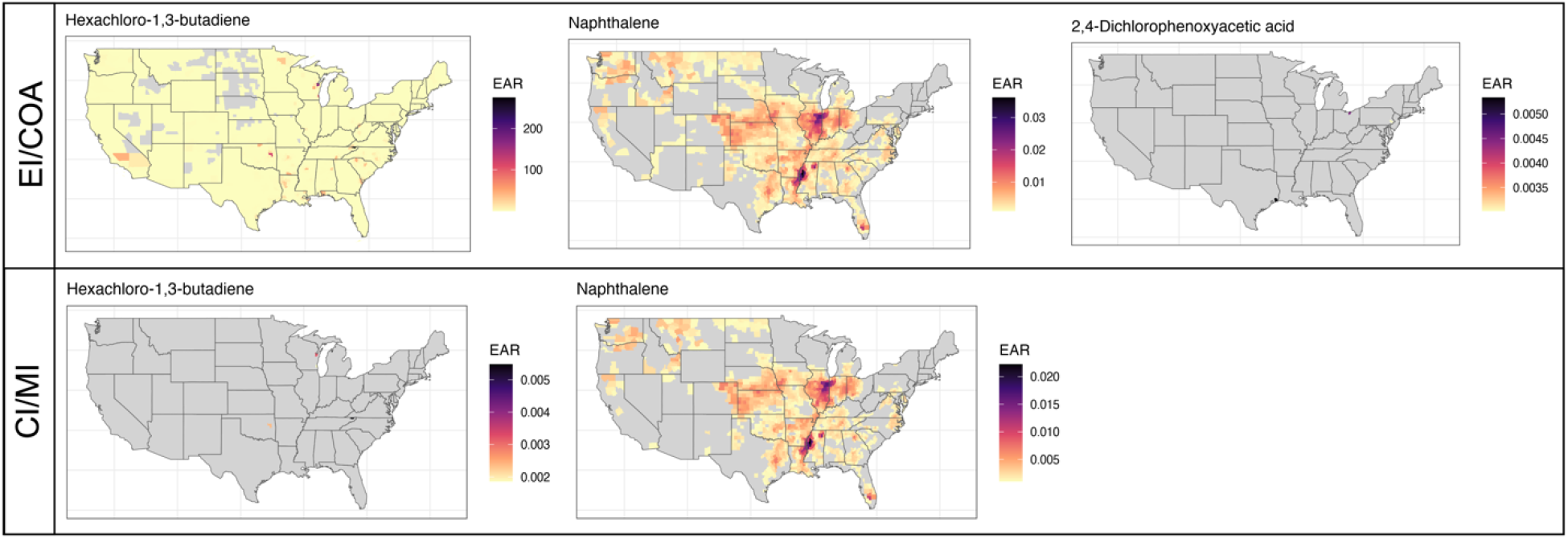
County-level maps of Exposure Activity Ratios (EAR) for NATA chemicals with activity in the endothelial injury/coagulation and cardiomyocyte/myocardial injury cardiovascular failure modes. EAR = county-level internal steady-state plasma concentration/most sensitive *in vitro* CV bioactivity concentration. Grey counties: EAR below the threshold of concern (0.001). Colored counties: EAR ≥ 0.001, with yellow/orange indicating lower and purple/black indicating higher EAR values.

## Discussion

This study demonstrates a multi-scale framework for understanding chemical cardiovascular effects, integrating *in vitro* HTS bioactivity data with reverse dosimetry modeling, human exposure predictions, and *in vivo* animal toxicity values for 859 chemicals, and geospatial risk projections for a subset with spatial distribution information. Such data-driven chemical assessments, anchored in human biology, and with uncertainty characterization, highlight the cardiovascular system as a sensitive and essential toxicological target, and optimize resource efficiency and translational relevance.

Here, we demonstrated that the minimum *in vitro* bioactivity concentration from a set of CV-relevant assays could serve as a threshold for potential *in vivo* toxicities, and that the BER is a flexible prioritization metric that can be calibrated by adjusting the choice of exposure estimate and uncertainty tolerance. The choice of PBPK model provides a range of extrapolated dose values that incorporate experimental and parameterization uncertainty, as do the human exposure predictions based off Bayesian probabilistic estimates. Here, the three models (1-C, 3-C, and PBTK models) produced broadly comparable EAD values, though systematic differences were observed. The 1C model, which assumes complete plasma availability (unbound fraction) and estimates hepatic clearance from unbound fraction to plasma protein, liver blood flow, and *in vitro* intrinsic clearance, consistently predicted higher EADs than the multi-compartment 3C and PBTK models. The 3C and PBTK models account for gut absorption, first-pass hepatic extraction, tissue partitioning, and organ-specific clearance, these processes predict lower steady-state plasma concentrations per unit dose and therefore generate lower EADs via reverse dosimetry, consistent with the protein-binding-dependent adjustment described above.

For 96.4% of the 695 chemicals with both *in vitro* and in vivo data, EAD_ivive_ values were equal to or lower than the POD_invivo_ from *in vivo* toxicology studies (positive POD ratio), affirming that CV-relevant *in vitro* endpoints are sensitive indicators of systemic toxicity. A subset of 88 chemicals showed both a positive POD ratio and a negative BER, identifying a group where the *in vitro* approach is simultaneously more conservative than animal data and where human exposure already fall within the bioactive range, compounds where adoption of *in vitro*-derived EADs as the primary screening metric would be most health-protective.

The 25 chemicals with a negative POD ratio, where *in vivo* animal studies identified hazard at lower doses than those extrapolated from the *in vitro* assays, share several common characteristics. They include organophosphate or organochlorine pesticides (including fenamiphos, parathion, phorate, heptachlor, lindane, DDT, dicrotophos, and azinphos-methyl) or per- and polyfluoroalkyl substances (PFAS) notably PFOS, whose primary toxicity mechanisms involve chronic neurotoxic, endocrine-disrupting, or bioaccumulative effects that unfold over prolonged exposure durations not captured by the *in vitro* protocols.

The range of BER values was substantially wider when using the upper 95th percentile ExpoCast estimate compared to the median, this is consistent with prior work demonstrating BER sensitivity to exposure metric choice ^38,47^. ExpoCast SEEM3 is a machine-learning based, probabilistic low-tier exposure assessment, and the 2 log10 units in the BER between median or 95th percentile exposure predictions reflect the broad credible intervals inherent in this approach. Of the 815 chemicals assessed, 12.52% had BER < 0, and 17 chemicals had a BER < −2, meaning potential human exposure exceeded the *in vitro*-derived bioactive dose by more than 100-fold. For these, the *in vivo*-derived MOE was also below −2, providing convergent evidence from both data streams for regulatory priority. Among the highest-priority chemicals, PFAS compounds, including PFOS and PFOA, are of particular concern given established epidemiological links between PFAS exposure and cardiovascular disease risk^51–54^.

The EAD_ivive_ reflects the dose required to perturb CV-relevant molecular targets, but may not be sufficient to produce adverse effects, as systemic toxicity is often multifactorial and compensatory biological pathways may intervene. Conversely, POD_invivo_ represents a threshold for observed adversity and reflects any systemic effect, not specifically cardiovascular perturbation. That >96% of chemicals nonetheless show a positive POD ratio strongly suggests that CV-relevant *in vitro* endpoints are sensitive indicators of broader systemic toxicity potential and may be more appropriate for chemical prioritization than non-specific *in vivo* PODs. Discrepancies for a small number of chemicals, such as DDT and Benzotrichloride, where *in vitro* methods predicted higher doses than *in vivo* studies, highlight limitations in capturing complex pharmacokinetics *in vitro*, particularly for chemicals whose toxicity unfolds over chronic exposure durations. Immune competent, multi-organoid platforms with the potential to capture the temporal progression from acute injury through persistent inflammatory remodeling are well-poised to more accurately model sustained chemical exposure scenarios and their cumulative impact on cardiovascular disease pathogenesis^55^.

The geospatial analysis revealed significant regional heterogeneity in CV chemical exposure risk across the United States. EAR values for chemicals such as naphthalene are concentrated in Illinois, Indiana, Arkansas, Mississippi, and Louisiana, regions associated with higher industrial activity and elevated CVD mortality rates (https://www.cdc.gov/nchs/pressroom/sosmap/heart_disease_mortality/heart_disease.htm), raising the possibility that co-exposure to CV-relevant environmental chemicals may contribute to the regional burden of cardiovascular disease. A key limitation is the uneven availability of detailed geospatial exposure data: while the NATA database provides robust coverage for air pollutants, comparable data are lacking for pesticides, plasticizers, and PFAS, several of which were flagged as high priority by the BER analysis. Expanding geospatial exposure modeling and improving population-level exposure models that account for demographic variability, including in toxicokinetics, would substantially strengthen the geospatial component of this framework^56^.

The findings reported here are timely in the context of an accelerating international regulatory shift toward NAM-based chemical safety evaluation. The FDA Modernization Act 2.0 (December 2022) removed the longstanding statutory mandate requiring animal testing for new drug applications, explicitly authorizing the use of *in vitro*, computational, and microphysiological systems as alternatives. This was followed in April 2025 by the FDA’s publication of a Roadmap to Reducing Animal Testing in Preclinical Safety Studies, which establishes a stepwise framework for replacing mandatory animal studies with human-relevant methods (https://www.fda.gov/news-events/press-announcements/fda-announces-plan-phase-out-animal-testing-requirement-monoclonal-antibodies-and-other-drugs). Concurrently, the NIH announced a major initiative to prioritize human-based research, including the planned creation of the Office of Research Innovation, Validation, and Application (ORIVA) to coordinate agency-wide NAM development, and a new policy requiring all NIH funding opportunities related to animal model systems to also support human-focused approaches such as NAMs (https://www.nih.gov/news-events/news-releases/nih-prioritize-human-based-research-technologies). For environmental chemical risk assessment specifically, the EPA’s ongoing New Approach Methods Work Plan and its January 2026 recommitment to eliminating mammalian animal testing by 2035 (https://www.pcrm.org/news/news-releases/physicians-committee-applauds-epa-recommitting-replacing-animal-tests) reflect growing regulatory expectation that HTS bioactivity data, IVIVE-derived EADs, and exposure-based metrics of the type generated here will play a central role in future chemical prioritization and risk management decisions. For highly prioritized chemicals, recent advances in engineered cardiac organoids and microphysiological systems hold tremendous promise as human-relevant platforms for functionally validating and characterizing chemicals identified through HTS-based screening, enabling mechanistic interrogation of cardiotoxicity and improving confidence in downstream risk assessment.^57^ The multiscale framework presented in this study, with its transparent, tiered metrics and reproducible computational workflow, is well-positioned to contribute to these evolving regulatory frameworks, particularly for the large number of environmental chemicals that currently lack adequate data concerning their cardiovascular effects.

## Conclusion

This study demonstrates that an integrated computational and human-based experimental framework provides a scalable and biologically-grounded approach for prioritizing environmental chemicals with potential cardiovascular risk. Across more than 800 environmental chemicals, pharmaceuticals, food additives, contaminants, and industrial compounds, human-relevant *in vitro* data revealed greater sensitivity than traditional animal toxicology in over 95% of cases, uncovering bioactivities at exposure-relevant levels that may be missed by conventional approaches. These findings underscore the cardiovascular system as a highly sensitive and underutilized indicator of potential chemical-induced adversity. By leveraging human-based NAMs, this framework enables more rapid, mechanistically informed, and human-relevant evaluation of cardiotoxicity, supporting proactive strategies for cardiovascular disease prevention and regulatory decision making.

## Non-standard Abbreviations & Acronyms

AC50: half-maximal activity concentration
ADMET: absorption, distribution, metabolism, and excretion–toxicity
BER: bioactivity exposure ratio
CPDat: EPA Chemical and Products Database
CV: cardiovascular
CVD: cardiovascular disease
EAD: equivalent administered dose
EAR: exposure activity ratio
EI/COA: endothelial injury/coagulation
FDA: US Food and Drug Administration
HTS: high-throughput screening
ICE: Integrated Chemical Environment
IVIVE: *in vitro* to *in vivo* extrapolation
IV: intravenous
MOE: margin of exposure
NAMs: new approach methodologies
NATA: National Air Toxics Assessment
NIH: National Institutes of Health
NOAEL: no observed adverse effect level
NOEL: no observed effect level
ORIVA: Office of Research Innovation, Validation, and Application
PBPK: physiologically based pharmacokinetics
PBTK: physiologically based toxicokinetic
PFAS: per- and polyfluoroalkyl substances
POD: point of departure
SEEM3: Systematic Empirical Evaluation of Models version 3
ToxValDB: The Toxicity Value Database
ΔAP: change in action potential
ΔINO: change in inotropy
ΔVA: change in vasoactivity

